# Genderized Gut and Oral Microbiome Shifts: Uncovering Sex-Specific Dysbiosis in Pancreatic Cancer

**DOI:** 10.1101/2024.10.02.616338

**Authors:** Zara Ahmed Khan, Mahin Ghorbani, Leon Heffinger, Anastasios Damdimopoulos, Carlos Fernández Moro, Mikael Björnstedt, J.-Matthias Löhr, Rainer Heuchel, Margaret Sällberg Chen, Dhifaf Sarhan

**Affiliations:** Department of Laboratory Medicine, Division of Pathology, Karolinska Institutet, Stockholm, Sweden; Bioinformatics and Expression Analysis Core Facility, Department of Biosciences, Karolinska University Hospital, Stockholm, Sweden; Nutrition, Karolinska Institute, Stockholm, Sweden, Department of Pathology and Cancer Diagnostics, Karolinska University Hospital, Stockholm, Sweden; Pancreatic cancer research lab, Department of Clinical Science, Intervention and Technology, Karolinska Institutet, Stockholm, Sweden

**Keywords:** Pancreatic ductal adenocarcinoma, microbiome, sex differences, dysbiosis, metagenomics

## Abstract

**Background:** Pancreatic ductal adenocarcinoma (PDAC) is one of the deadliest cancers, responsible for approximately 466,000 deaths globally in 2020. Its incidence increases by about 1% annually, with a higher occurrence in males than females. While differences in immune responses and tumor biology between sexes have been explored, the role of the microbiome in gender-specific PDAC progression is still unclear. Investigating these differences could offer crucial insights for personalized treatment strategies for males and females.

**Methods:** This study reanalyzed oral and gut microbiome data from BioProject: PRJNA832909, comprising 191 samples from PDAC patients and healthy controls. Using shotgun metagenomic sequencing, we examined gender-specific bacterial signatures. Alpha diversity (richness) and beta diversity (community composition) were analyzed. Differentially abundant bacterial taxa were identified via LEfSe, and gender-specific bacterial panels were validated using CombiROC.

**Results:** Alpha diversity analysis revealed significant differences in microbial richness, particularly between male and female PDAC patients and their healthy controls. Beta diversity demonstrated distinct microbial shifts between the PDAC and control groups across genders. LEfSe identified several pathogenic bacteria contributing to gender-specific dysbiosis, including *Streptococcus, Fusobacterium*, and *Prevotella*. Shared and sex-specific bacterial species in PDAC were highlighted through Venn diagram analysis. CombiROC validated the predictive ability of these bacterial markers, with AUC values exceeding 0.90 for both sexes.

**Conclusion:** This study uncovered gender-specific microbial patterns in PDAC patients, potentially influenced by sex-specific immune responses. These findings provide important insights into the progression of PDAC and support sex-targeted diagnostic and therapeutic interventions.

**Graphical abstract:** 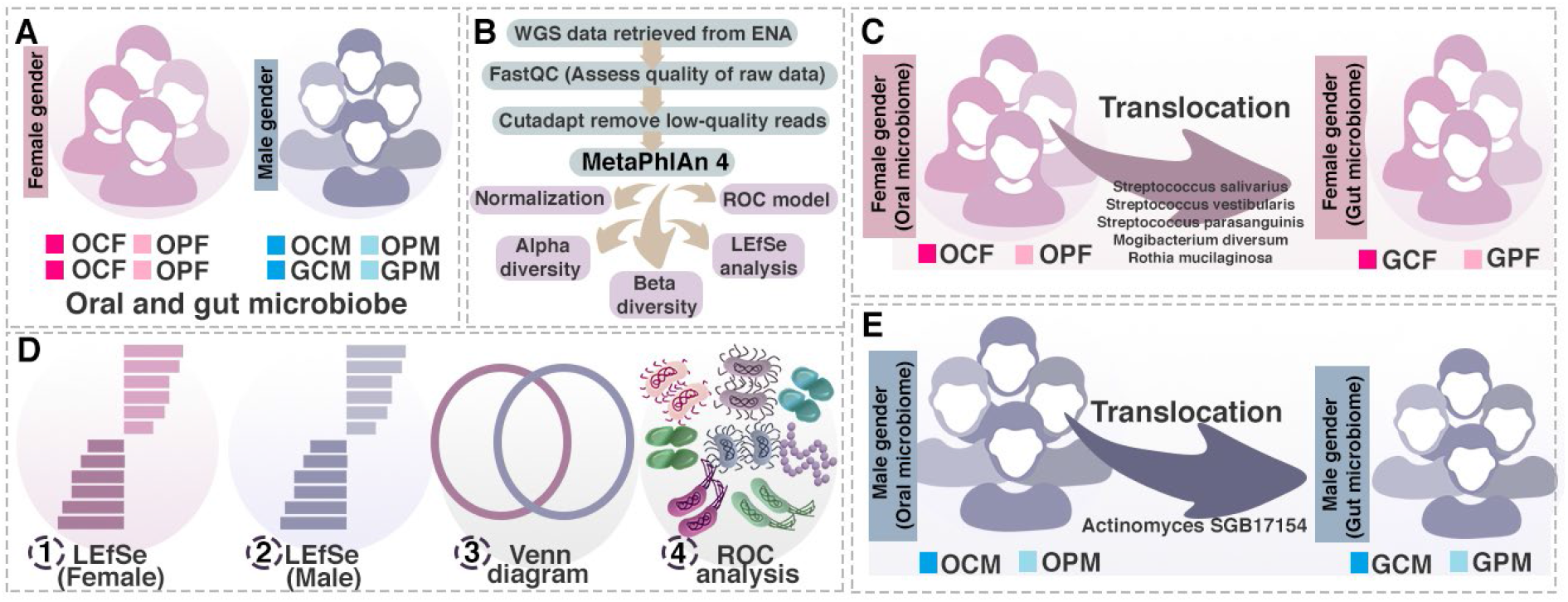

## Introduction

Pancreatic cancer (PC) presents an enormous therapeutic challenge with its highly immunosuppressive tumor microenvironment (TME), contributing to the poor 12% 5-year overall survival rate.^1^ Even though prognosis slightly improved over recent years, there is still an unmet need to further improve and develop therapy options. Therefore, a deeper understanding of the PC-TME is urgently needed. Geller et al. investigated the potential influence of intratumor bacteria in mediating chemotherapeutic resistance. This underscores the importance of exploring the complex interplay between the microbiome and tumor biology in the TME of PC^2^. Around 90% of PC cases arise from pancreatic ductal cells resulting in pancreatic ductal adenocarcinoma (PDAC) being the most common PC type^3^. Due to its anatomical proximity to the duodenum, the PDAC-TME is hypothesized to be influenced by retrograde infiltration of microbiota from the gastrointestinal (GI) tract^2^. Recent studies have linked variations in bacterial signatures within the tumor microenvironment to patient survival, positioning the tumor-associated microbiome as a focal point for current research efforts^4^. We and others have detected several bacteria species in PDAC TME^5, 6, 7^. Specific bacterial species and peptides have been detected in PDAC tissues, with evidence suggesting that these microbial components may directly interact with immune cells, thereby impacting tumor progression and response to treatment^8^. Notably, sex-based differences in immune responses and cancer progression have gained increasing recognition, yet their influence on the PDAC-associated microbiome and subsequent tumor biology has not been thoroughly investigated. Our recent work identified the macrophage receptor Formyl-peptide receptor 2 (FPR2) as a critical mediator of PDAC progression in female mice, highlighting the potential for sex-specific immune pathways to shape the TME^9^. However, the role of the microbiome in mediating these sex differences remains unclear. Given that the GI tract and oral cavity harbor distinct microbial communities that influence not only local immunity but also systemic disease processes, investigating sex-specific differences in these microbial populations could provide novel insights into PDAC pathogenesis. Gonadal steroid hormones, known to modulate both the immune system and microbiome composition, may further contribute to sex-dependent variations in disease outcomes. Thus, delineating the distinct microbial signatures in male and female PDAC patients could uncover new mechanisms underlying these disparities and suggest tailored therapeutic strategies.

In this study, we aim to characterize the sex-specific gut and oral microbial profiles in PDAC patients to identify sex-biased variations that may influence tumor progression and therapeutic response. By elucidating these microbiome-driven differences, we seek to establish a foundation for personalized, sex-specific treatment approaches, ultimately enhancing therapeutic efficacy and improving patient outcomes in PDAC.

## Methodology

### Sample Retrieval and Study Design

In this study, we utilized data from Project: PRJNA832909, which investigated the metagenomics of the fecal and salivary microbiome in pancreatic cancer patients, focusing on identifying sex-specific bacteria contributing to pancreatic cancer pathogenesis. Our analysis incorporated 191 samples from a cohort referenced in the study *Metagenomic Identification of Microbial Signatures Predicting Pancreatic Cancer from a Multinational Study* (Nagata et al., 2022). These samples included 28 Gut Control Female (GCF), 21 Gut PDAC Female (GPF), 30 Gut Control Male (GCM), 19 Gut PDAC Male (GPM), 28 Oral Control Female (OCF), 17 Oral PDAC Female (OPF), 31 Oral Control Male (OCM), and 17 Oral PDAC Male (OPM).

### Bioinformatics and Statistical Analysis

In our study, raw sequencing reads were quality-checked using FastQC v0.11.8 to assess base quality and sequence integrity, followed by adapter trimming with Cutadapt v2.8 to remove extraneous sequences. Taxonomic profiling was conducted using MetaPhlAn4, which utilizes clade-specific marker genes from over 17,000 reference genomes encompassing bacterial, archaeal, viral, and eukaryotic taxa. Taxonomic assignments were made with the in-house MetaPhlAn4 database, and feature counts were filtered and normalized using Total Sum Scaling (TSS) to standardize the data for downstream analyses.

Alpha diversity metrics, specifically Observed Richness and the Shannon Index, were calculated to quantify species richness and evenness, respectively. Beta diversity, representing inter-sample community variation, was evaluated using Bray-Curtis and Jaccard distances and visualized via Nonmetric Multidimensional Scaling (NMDS). Statistical significance for beta diversity comparisons was assessed using Permutational Multivariate Analysis of Variance (PERMANOVA) with a threshold of P < 0.05.

For differential abundance analysis, we utilized LEfSe v1.1.01 (Linear Discriminant Analysis Effect Size) with an LDA score threshold >2 and P < 0.05 to identify taxa significantly enriched between female control versus female PDAC samples, and male control versus male PDAC samples, across both the oral and gut microbiomes. The identified bacterial markers were subsequently validated using CombiROC, a tool for combining high-performing metagenomic features from LEfSe to enhance classification performance. The resulting sex-specific bacterial panels were subjected to independent post-hoc Receiver Operating Characteristic (ROC) curve analysis (P < 0.05) for additional validation (see Supplementary File). This integrative analytical framework allowed us to establish robust, sex-specific bacterial markers for distinguishing pancreatic cancer in male and female cohorts.

### Methods titration IF

To validate the presence of sex-specific bacterial strains and their interactions with tumor-associated immune cells, paraffin-embedded pancreatic tumor tissues were subjected to spatial analysis. Tissue sections were prepared by initial overnight incubation at 37°C, followed by an additional hour at 60°C. Rehydration was achieved through a series of washes: three consecutive washes in Xylene (15 minutes each), followed by three washes in 100% Ethanol (15 minutes each), two washes in 95% Ethanol (15 minutes each), a single wash in 70% Ethanol, and finally a 10-minute incubation in Phosphate-Buffered Saline (PBS).

Antigen retrieval was conducted using a 1:100 dilution of Vector Antigen Unmasking Solution (Cat# H3300) with microwave heating at 800 W for 20 minutes, followed by a 20-minute cooling period at room temperature. Tissues were then blocked using 5% goat serum in PBS to minimize non-specific binding. Primary antibody staining was performed using different dilutions (1:10, 1:50, and 1:100) of antibodies targeting Lipopolysaccharide Core (Cat# HM6011-20UG) and Lipoteichoic Acid (Cat# HM5018-20UG) for one hour in a humidified chamber. A PBS-treated slide was used as a negative control.

Following primary antibody incubation, slides were washed in PBS for 5 minutes and then incubated with an AF488-conjugated secondary antibody (Cat# A11001) at a 1:1000 dilution for 30 minutes. After a subsequent PBS wash (5 minutes), slides were counterstained with DAPI at a 1:2500 dilution for 10 minutes. The slides underwent two additional PBS washes (5 minutes each) before being mounted with an anti-quenching mounting media (Cat# P36961). Imaging was performed using the Vectra 3 Automated Quantitative Pathology Imaging System. Background subtraction and image processing were conducted using inForm 2.6.0 software to ensure precise quantification and visualization of the bacterial and immune cell interactions within the tumor microenvironment.

## Results

### Sex-Specific Analysis Design

The study by Nagata et al. provided critical insights into the gut and oral microbiomes of PDAC patients, identifying specific microbial signatures predictive of pancreatic cancer. Key findings included the enrichment of *Streptococcus* and *Veillonella* species and a marked depletion of *Faecalibacterium prausnitzii*, which collectively demonstrated high predictive accuracy for PDAC across diverse international cohorts. By rigorously controlling for confounding variables such as age, sex, body mass index (BMI), and lifestyle factors (e.g., smoking and alcohol consumption), the study ensured that observed microbiome differences were primarily attributable to disease status.

Building on these findings, our study conducted a detailed, sex-specific analysis to explore microbial variations between male and female PDAC patients. We compared the oral and gut microbiome profiles of female PDAC patients to those of healthy female controls to identify bacterial taxa specifically associated with PDAC in women. Similarly, we analyzed the microbiome composition in male PDAC patients versus healthy male controls to uncover bacterial species uniquely contributing to PDAC pathogenesis in men.

Following these sex-specific analyses, we employed a Venn diagram approach to categorize bacteria shared between both sexes and those unique to either males or females. This strategy enabled us to define common microbial signatures associated with PDAC while simultaneously highlighting specific bacterial profiles, thereby enhancing our understanding of sex-biased differences in microbiome dysbiosis and their role in pancreatic cancer development. Our findings are consistent with the growing body of evidence emphasizing the importance of sex as a biological variable in both microbiome research and cancer progression.

### Microbiome Composition of oral microbiome at phylum, genus, and species level

At the phylum level, the oral microbiomes of PDAC patients and healthy controls were primarily dominated by *Firmicutes, Actinobacteria, Bacteroidetes*, and *Proteobacteria* in both sexes. However, notable sex-specific differences were observed. Female PDAC patients exhibited a significant increase in *Bacteroidetes* and a concomitant decrease in *Actinobacteria* relative to healthy female controls. In male PDAC patients, *Bacteroidetes* levels were also elevated compared to healthy males, though to a lesser extent than in females. Interestingly, *Proteobacteria* levels showed opposing trends between sexes: male PDAC patients had higher *Proteobacteria* levels compared to healthy males, while female PDAC patients exhibited lower *Proteobacteria* levels relative to healthy females (**Figure 1A**). These cross-sex comparisons highlight both shared and distinct microbiome shifts at the phylum level in male and female PDAC patients.

**Figure 1.**
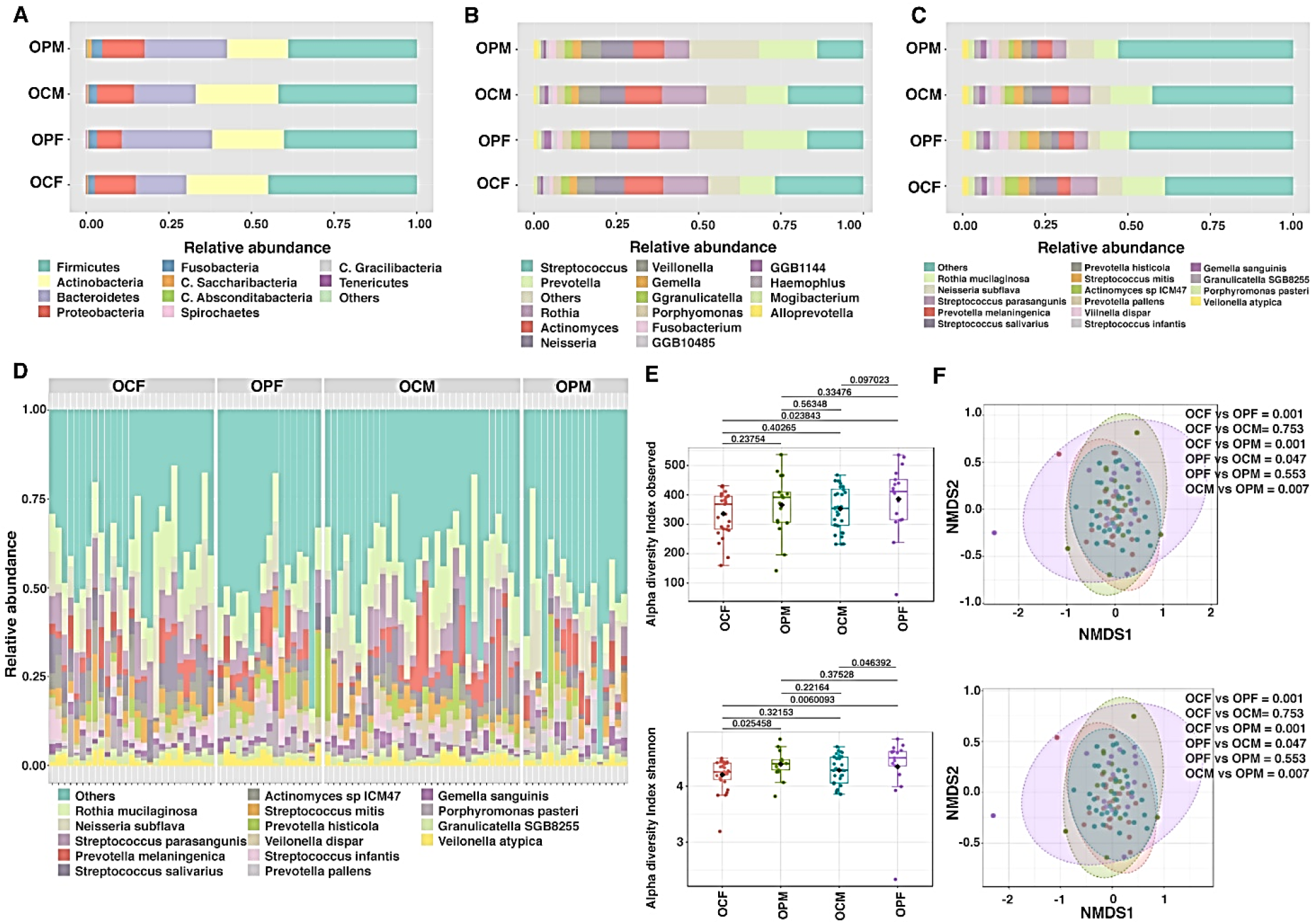
Oral Microbiome Composition, Diversity, and Variation Between PDAC and Control Groups by Sex. (A) Microbiome Composition: This panel shows the relative abundance of the oral microbiome at the phylum level across four groups: oral control females (OCF), oral PDAC females (OPF), oral control males (OCM), and oral PDAC males (OPM). Dominant phyla include *Firmicutes, Actinobacteria, Bacteroidetes, and Proteobacteria*.(B) Genus-Level Composition: Relative abundance of bacterial genera across the same groups. Genera such *as Streptococcus, Prevotella*, and *Rothia* dominate in the oral microbiomes, with variations between PDAC patients and controls. (C) Species-Level Composition: Relative abundance of bacterial species in the four groups. Species like *Streptococcus salivarius* and *Rothia mucilaginos*a are among the top species, with clear differences observed between sexes and PDAC statuses. (D) Relative Abundance at Species Level in individual samples visualization for Top 15 Bacteria: Stacked bar plot showing the species-level composition for the top 15 most abundant species, illustrating differences between controls and PDAC patients in both sexes. (E) Alpha Diversity: Boxplots displaying the Observed and Shannon indices, which measure species richness and diversity. Significant differences in alpha diversity between PDAC patients and controls are annotated with p-values. (F) Beta Diversity: Non-metric multidimensional scaling (NMDS) plots showing beta diversity using Bray-Curtis dissimilarity and Jaccard index, visualizing distinct microbial community compositions across groups. Significant differences between groups are validated by PERMANOVA tests (p < 0.05).

At the genus level, differences in microbial composition were further delineated. In female PDAC patients, a marked decrease in *Streptococcus* and a significant increase in *Prevotella* were observed when compared to healthy female controls. A similar trend was noted in male PDAC patients, although the relative abundance of these genera varied between sexes. Consistent across both sexes was the reduction of *Rothia* and *Actinomyces*. Notably, *Neisseria* decreased significantly in female PDAC patients but showed a slight increase in male PDAC patients compared to their respective controls. Additionally, species such as *GGB1144* exhibited distinct sex-specific patterns (**Figure 1B**), indicating that although certain microbial changes were shared, there were also unique shifts in the composition of oral microbiomes between male and female PDAC patients, which could play a role in differential disease pathogenesis.

At the species level, the oral microbiomes of PDAC patients and healthy controls were dominated by *Rothia mucilaginosa, Neisseria subflava, Streptococcus parasanguinis, Prevotella melaninogenica*, and *Streptococcus salivarius* across both sexes. However, specific alterations in species abundance were evident in PDAC patients. In both male and female PDAC cohorts, there was a significant reduction in *Streptococcus salivarius* and *Rothia mucilaginosa* relative to healthy controls. In females, *Prevotella melaninogenica* showed a pronounced increase in PDAC patients compared to healthy females, whereas this increase was less substantial in males. Furthermore, *Neisseria subflava* displayed an opposite trend between the sexes, with decreased abundance in female PDAC patients and a slight increase in male patients (**Figure 1C, Figure 1D**), underscoring distinct, sex-specific dysbiosis patterns at the species level. These findings suggest that sex-specific microbiome alterations may contribute to the differential pathogenesis of pancreatic cancer in male and female patients.

### Alpha and beta diversities of oral microbiome

Significant differences in microbial richness and diversity were observed between PDAC patients and healthy controls, with notable variations evident in sex-specific comparisons. Alpha diversity analysis, which assesses intra-sample diversity, revealed no major differences except a significant difference observed in species richness between oral microbiomes of Oral Control Female (OCF) and Oral PDAC Male (OPM) (P = 0.023). Indeed, Shannon diversity indicated significant variations between female controls and female PDAC patients (P = 0.025), female controls and male PDAC patients (P = 0.006), and male controls versus male PDAC patients (P = 0.046), suggesting substantial sex-specific shifts in microbial diversity (**Figure 1E**).

Beta diversity analysis, using both Bray-Curtis and Jaccard indices, further underscored these distinctions by capturing inter-sample diversity and compositional dissimilarities. Significant separation was observed between female controls and female PDAC patients (Bray-Curtis: p = 0.001; Jaccard: p = 0.001), female controls and male PDAC patients (Bray-Curtis: p = 0.001; Jaccard: p = 0.001), and male controls versus male PDAC patients (Bray-Curtis: p = 0.007; Jaccard: p = 0.007). Additionally, the Jaccard index detected significant differences between female and male PDAC patients (p = 0.042), indicating sex-specific community composition shifts (**Figure 1F**).

These findings collectively highlight pronounced sex-specific microbial dysbiosis patterns at both the alpha and beta diversity levels, suggesting that the oral microbiome may play a differential role in PDAC pathogenesis based on sex. Such distinctions emphasize the need for sex-stratified analyses in microbiome studies of pancreatic cancer to better elucidate the underlying microbial contributions to disease progression.

### Microbiome Composition of gut microbiome at phylum, genus, and species level

Next, we investigated the microbiome of the gut. At the genus level, the gut microbiomes of PDAC patients and healthy controls are characterized by the predominance of *Bacteroides, Phocaeicola, Blautia, Bifidobacterium, Faecalibacterium*, and *Prevotella* in both sexes. However, female PDAC patients exhibited a significant increase in *Prevotella* relative to healthy females, while this genus showed minimal variation between male PDAC patients and their healthy male counterparts, suggesting a potential sex-specific role for *Prevotella* in PDAC pathogenesis. Similarly, *Bifidobacterium* displayed divergent abundance patterns between male and female PDAC patients compared to their respective controls. Although many genera, such as *Ruminococcus* and *Streptococcus*, demonstrated consistent shifts in both sexes, these results indicate that while some microbial alterations in PDAC are shared, others may be sex-specific (**Figure 2B**).

**Figure 2.**
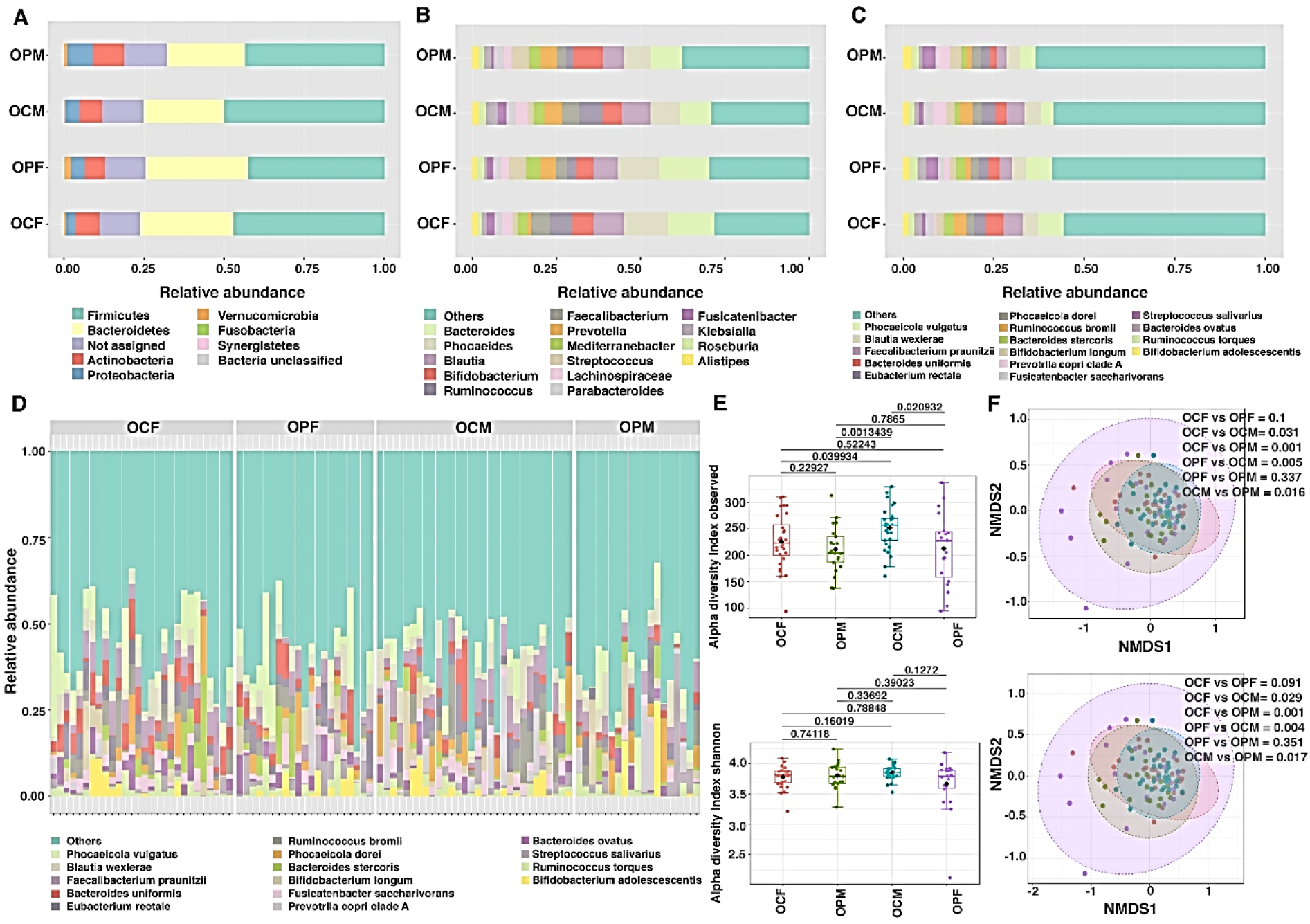
Gut Microbiome Composition, Diversity, and Variation Between PDAC and Control Groups by Sex. (A) Microbiome Composition: This panel displays the relative abundance of the gut microbiome at the phylum level for female controls (OCF), female PDAC patients (OPF), male controls (OCM), and male PDAC patients (OPM). Dominant phyla include *Firmicutes, Bacteroidetes, Actinobacteria, and Proteobacteria*. (B) Genus-Level Composition: Relative abundance of bacterial genera across the same groups, highlighting top dominant genera (C) Species-Level Composition: Relative abundance of top dominant bacterial species in the four groups show variations between PDAC patients and controls across sexes. (D) Relative Abundance at Individual Sample Visualization of Species Level for Top 15 Bacteria: The bar plot shows the top 15 most abundant species in the gut microbiomes of all groups, highlighting differences in microbial composition between controls and PDAC patients. (E) Alpha Diversity: Boxplots for alpha diversity metrics (Observed index and Shannon index) show the differences in microbial richness and diversity between PDAC patients and controls, with significant sexes-specific trends annotated by p-values. (F) Beta Diversity: NMDS plots visualize beta diversity using Bray-Curtis dissimilarity and Jaccard index, showing distinct microbial community compositions between groups. PERMANOVA test results confirm significant beta diversity differences between subgroups (p < 0.05).

At the species level, the top 15 species dominating the gut microbiomes of PDAC patients and healthy controls included *Phocaeicola dorei, Blautia wexlerae, Faecalibacterium prausnitzii*, and *Bifidobacterium longum* across both sexes. While the overall trend in abundance shifts for these species was largely similar in male and female PDAC patients, certain subtle differences were observed. For instance, *Fusciantenibacter saccharivorans* showed a higher abundance in healthy control males compared to male PDAC patients, but no significant change was noted in females. Although more than 50% of species in the gut microbiome were not among the top 15 identified species, these findings suggest a combination of shared and sex-specific microbial shifts at the species level in PDAC patients (**Figure 2C and Figure 2D**).

### Alpha and Beta Diversity Analysis

Alpha diversity analysis revealed significant differences in microbial richness between PDAC patients and healthy controls, particularly in sex-stratified comparisons. Observed richness was significantly different between female and male healthy controls (p = 0.039), male healthy controls and male PDAC patients (p = 0.021), and female healthy controls and male healthy controls (p = 0.001). However, Shannon diversity did not show any statistically significant variations, suggesting that while species richness differed between groups, overall diversity and evenness were comparable (**Figure 2E**).

Beta diversity analysis, utilizing Bray-Curtis and Jaccard indices, demonstrated significant dissimilarities between sex and disease groups. Notably, differences were observed between female and male healthy controls (Bray-Curtis: p = 0.031; Jaccard: p = 0.029), female healthy controls and male PDAC patients (Bray-Curtis: p = 0.001; Jaccard: p = 0.001), female PDAC patients and male healthy controls (Bray-Curtis: p = 0.005; Jaccard: p = 0.004), and male healthy controls versus male PDAC patients (Bray-Curtis: p = 0.016; Jaccard: p = 0.017) (**Figure 2F**). These results highlight pronounced sex-specific microbial shifts in the gut microbiome of PDAC patients compared to healthy individuals.

Overall, our findings underscore the existence of both shared and distinct microbial dysbiosis patterns at various taxonomic levels, with clear sex-specific differences in both alpha and beta diversity.

### Discriminative Oral Bacterial Signature Identification and Sex-specific Predictive Model Development for Pancreatic Cancer Using LEfSe, Venn Analysis, and CombiROC

In our study, LEfSe analysis was used to identify significantly different bacteria between the gut microbiomes of female controls and female PDAC patients, as well as between male controls and male PDAC patients **(Figure 3B)**. A Venn diagram **(Figure 3C)** was then created to distinguish shared and sex-specific bacterial taxa. The analysis identified 25 species uniquely associated with females and 15 species specific to males, while 9 species were found to be shared between both sexes.

**Fig. 3.**
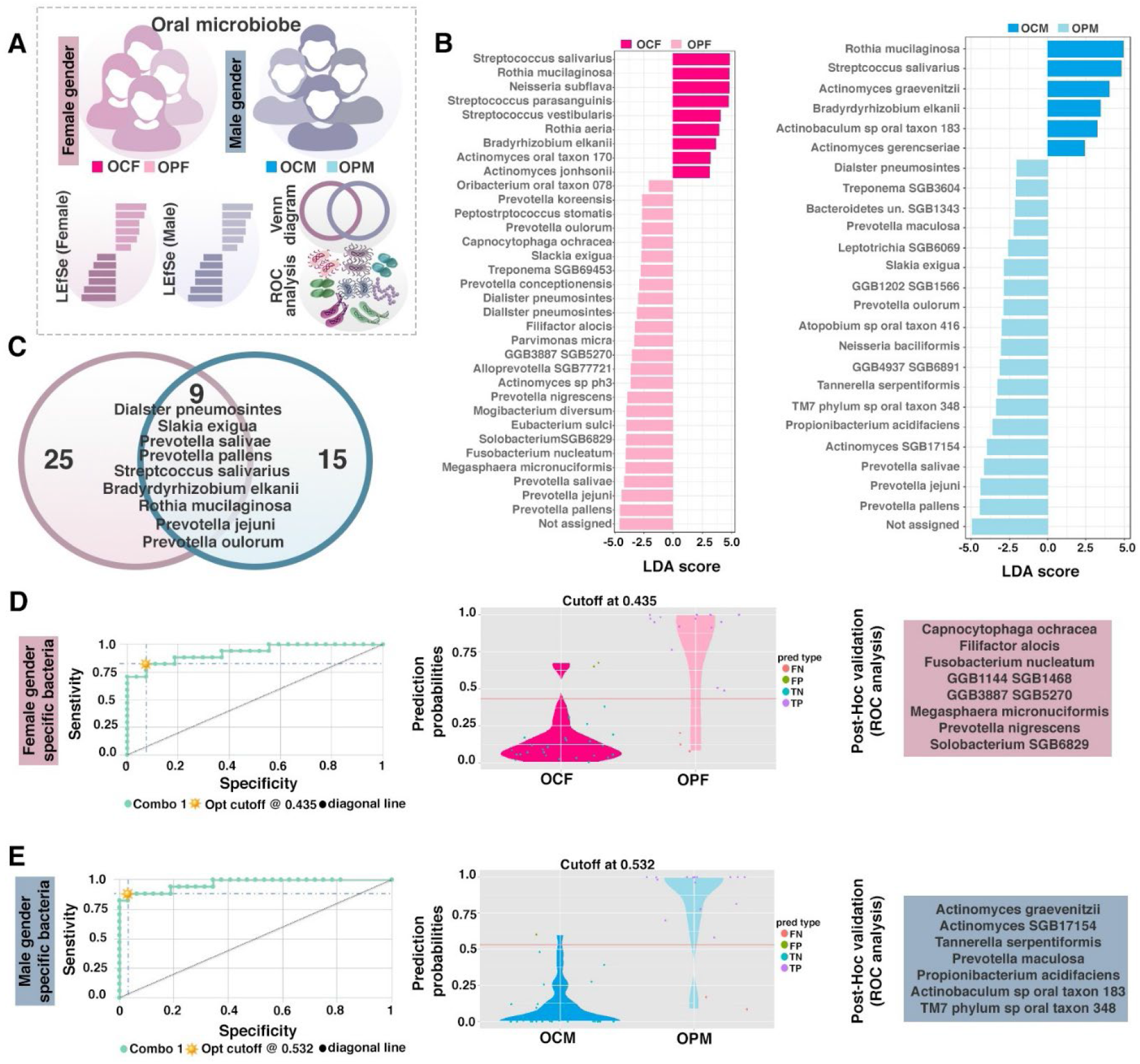
Sex-Specific Discriminant Bacterial Signatures in the Oral Microbiome of PDAC Patients and Healthy Controls. (B left) LEfSe Analysis for Female Oral Microbiota: Linear discriminant analysis effect size (LEfSe) plots showing the most discriminant bacterial species between oral control females (OCF) and oral PDAC females (OPF). The LDA scores highlight the species most significantly associated with PDAC or control groups in females. (B right) LEfSe Analysis for Male Oral Microbiota: LEfSe plots for male oral control (OCM) versus male PDAC patients (OPM). The LDA scores highlight the bacterial species that most significantly differentiate male PDAC patients from their healthy male counterparts. (C) Shared and Sex-Specific Oral Microbial Signatures: Venn diagram showing the overlap of bacterial species between the female PDAC bacterial panel (pink) and the male PDAC bacterial panel (blue). A total of 9 bacterial species are shared across sexes, while 25 species are specific to females and 15 species are specific to males, indicating sex-specific differences in the oral microbiome of PDAC patients compared to controls. (D) ROC Analysis for Female Oral Microbial Panel: Receiver operating characteristic (ROC) curve demonstrates the sensitivity and specificity of the female-specific bacterial markers identified from the oral microbiome for predicting PDAC in females. The AUC value of 0.92 (p < 0.05) indicates strong predictive power. The top significant bacteria of female-specific, selected by CombiROC software, are shown in Figure E. The violin plot visualizes the distribution of false positives, false negatives, true negatives, and true positives in the samples, with minimal false results.(E) ROC Analysis for Male Oral Microbial Panel: ROC curve for male-specific bacterial markers showing the performance of the predictive model for identifying PDAC in males. The AUC value of 0.96 (p < 0.05) indicates high predictive accuracy. The top significant bacteria of male-specific, listed in Figure E, were selected through CombiROC analysis. The violin plot demonstrates the sample distribution of false positives, false negatives, true negatives, and true positives, with a minimized number of false classifications.

To further evaluate the predictive power of the identified bacterial species, we employed CombiROC to determine the most significant bacterial panels for both females (**Figure 3D**) and males (**Figure 3E**). The resulting Receiver Operating Characteristic (ROC) curve analyses yielded Area Under the Curve (AUC) values of 0.92 for females and 0.96 for males, indicating robust predictive accuracy.

In female PDAC patients, the key bacterial species identified included *Capnocytophaga ochracea, Filifactor alocis, Fusobacterium nucleatum, GGB1144 SGB1468, GGB3887 SGB5270, Megasphaera micronuciformis, Prevotella nigrescens*, and *Solobacterium SGB6829*. These species underscore distinct sex-specific microbial dysbiosis patterns in pancreatic cancer, providing valuable insights for the development of targeted diagnostic and therapeutic strategies.

In male PDAC patients, the key bacterial species identified included *Actinomyces graevenitzii, Actinomyces SGB17154, Tannerella serpentiformis, Prevotella maculosa, Propionibacterium acidifaciens, Actinobaculum sp. oral taxon 183*, and *TM7 phylum sp. oral taxon 348*. These sex-specific microbial signatures provide critical insights into the unique microbiome dynamics in male PDAC patients, offering potential avenues for the development of personalized diagnostics.

The species shared between both sexes included *Dialister pneumosintes, Slackia exigua, Prevotella salivae, Prevotella pallens, Prevotella jejuni, Prevotella oulorum, Streptococcus salivarius, Bradyrhizobium elkanii*, and *Rothia mucilaginosa*, suggesting common microbial dysbiosis patterns that may contribute to pancreatic cancer pathogenesis across both sexes.

### Discriminative Gut Bacterial Signature Identification and Sex-Specific Predictive Model Development for Pancreatic Cancer Using LEfSe, Venn Analysis, and CombiROC

In our study, LEfSe analysis was employed to identify significantly different bacterial taxa in the gut microbiomes of female and male PDAC patients compared to their respective healthy controls. A Venn diagram was used to categorize shared and sex-specific bacterial species, revealing 27 species unique to females, 38 species unique to males, and 5 species common to both sexes.

To further assess the predictive power of these identified bacterial species, CombiROC software was utilized to construct optimal microbial panels for both male and female cohorts. The resulting ROC curve analyses demonstrated strong predictive accuracy, with AUC values exceeding 0.95 for females and 0.88 for males.

In female PDAC patients, the most significant bacterial species identified were Butyricimonas *SGB15260, GGB9712 SGB15244, Bacteroides uniformis, Mogibacterium diversum, Rothia mucilaginosa, Streptococcus oralis, Streptococcus parasanguinis, Streptococcus salivarius*, and *Sutterella wadsworthensis*. Conversely, in male PDAC patients, the key bacterial markers included *Agathobaculum butyriciproducens, Alistipes putredinis, Dorea longicatena, Fusicatenibacter saccharivorans, Lachnospira eligens, Lawsonibacter asaccharolyticus*, and *Sanguibacter SGB15121*.

Species shared between both sexes comprised *Streptococcus constellatus, Veillonella atypica, Veillonella parvula, Clostridiaceae bacterium*, and *GGB3746 SGB5089*, indicating a set of core microbial dysbiosis patterns potentially contributing to PDAC pathogenesis in both sexes.

### Shared Bacterial Species Between Oral and Gut Microbiomes in PDAC Patients: Insights into Sex-Specific Dysbiosis and Translocation Hypotheses

Next, we analyzed the shared bacterial species between the oral and gut microbiomes of female PDAC patients and healthy female controls (FF panel) using LEfSe analysis. By comparing the oral FF panel with the gut FF panel, we found five common species across both environments: *Mogibacterium diversum, Streptococcus salivarius, Streptococcus parasanguinis, Rothia mucilaginosa*, and *Streptococcus vestibularis* **(Figure 5B)**.

**Fig. 4.**
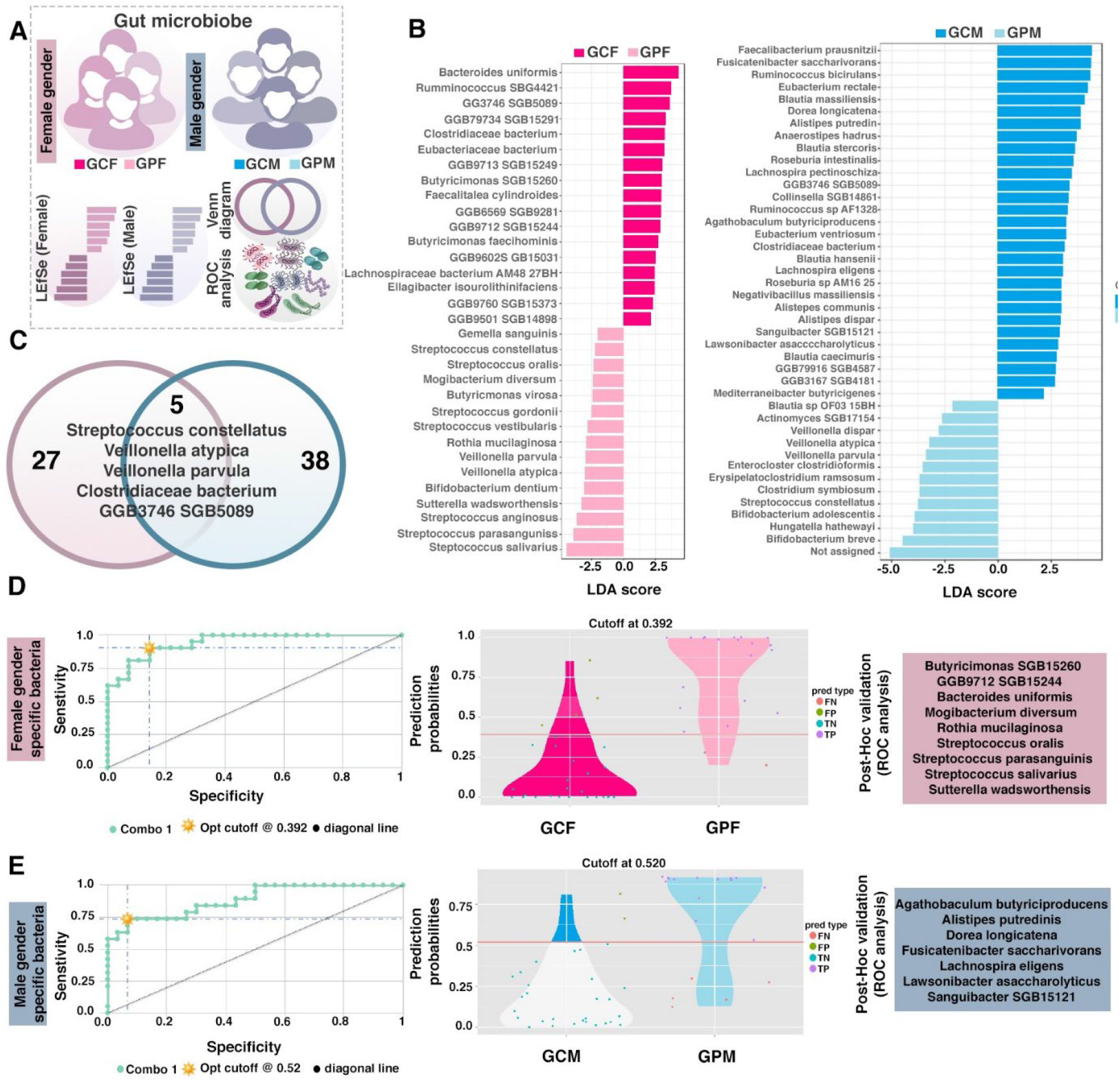
Comparison of Sex-Specific Gut Microbial Signatures in PDAC Patients vs. Healthy Controls. (A) LEfSe Analysis for Female Gut Microbiota: Linear discriminant analysis effect size (LEfSe) plot showing the bacterial species that most significantly differentiate between gut control females (GCF) and gut PDAC females (GPF). The LDA scores highlight the species most significantly associated with PDAC or control groups (B) LEfSe Analysis for Male Gut Microbiota: LEfSe plot comparing gut control males (GCM) and gut PDAC males (GPM), identifying the bacterial species that most significantly differentiate male PDAC patients from their healthy male counterparts. The LDA scores reflect the microbial differences specific to male PDAC patients. (C) Shared and Sex-Specific Gut Microbial Signatures: Venn diagram depicting the overlap of gut bacterial species between female PDAC patients (pink) and male PDAC patients (blue). A total of 5 species are shared between sexes, while 27 species are specific to females and 38 species are unique to males, indicating sex-specific microbiome differences in PDAC patients.(D) ROC Analysis for Female Gut Microbial Panel: Receiver operating characteristic (ROC) curve illustrating the sensitivity and specificity of female-specific bacterial markers for distinguishing PDAC patients from healthy controls and the violin plot shows the predicted probabilities for control (GCF) and PDAC (GPF) groups, with minimal false results. CombiROC software was used to identify the most significant bacterial panels, and the area under the curve (AUC) value for the female panel is above 0.95, demonstrating strong predictive power. (E) ROC Analysis for Male Gut Microbial Panel: ROC curve illustrating the performance of male-specific bacterial markers for predicting PDAC in males and the violin plot demonstrates the predicted probabilities for control (GCM) and PDAC (GPM) groups. The bacterial panel identified by CombiROC for males shows an AUC value of 0.88, indicating high predictive accuracy. and the violin plot shows the predicted probabilities for control (GCF) and PDAC (GPF) groups, with minimal false results

**Fig. 5.**
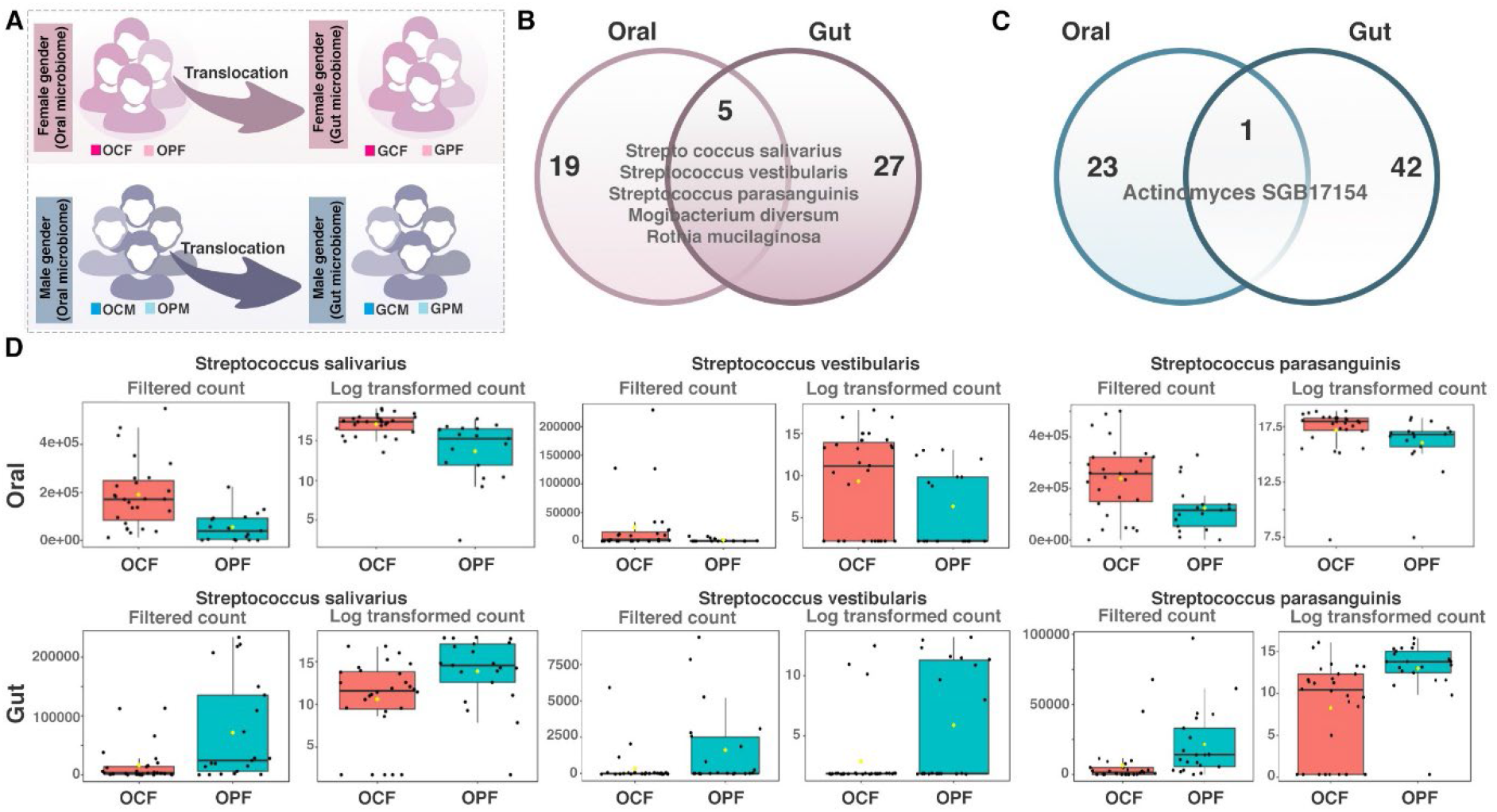
Shared Bacterial Species Between Oral and Gut Microbiomes in Female and Male PDAC Patients and Controls. (B) Shared Bacterial Species in Female Patients and Controls: Venn diagram showing the overlap of bacterial species identified by LEfSe between the oral and gut microbiomes in both female PDAC patients and healthy controls. Five bacterial species are common across both environments: *Streptococcus salivarius, Streptococcus vestibularis, Streptococcus parasanguinis, Mogibacterium diversum*, and *Rothia mucilaginosa*. These species demonstrate distinct enrichment patterns in the oral and gut microbiomes of female PDAC patients compared to controls. (C) Shared Bacterial Species in Male Patients and Controls: Venn diagram showing the overlap of bacterial species identified by LEfSe between the oral and gut microbiomes in both male PDAC patients and healthy controls. One bacterial species, *Actinomyces SGB17154*, is shared between the oral and gut microbiomes in males, suggesting a potential link in sex-specific microbial dysbiosis in PDAC patients and controls. (D) Abundance of Shared Bacterial Species in Females and Males (Oral and Gut): Box plots comparing the abundance of *Streptococcus salivarius, Streptococcus vestibularis*, and *Streptococcus parasanguinis* between oral control females (OCF) and oral PDAC females (OPF), as well as their corresponding gut samples. The top row shows the oral microbiome, and the bottom row shows the gut microbiome, with both filtered and log-transformed counts. These box plots illustrate the changes in abundance between controls and PDAC patients across both the oral and gut microbiomes.

Interestingly, while these species were detected in both the oral and gut microbiomes, they exhibited distinct patterns of enrichment. In healthy individuals, these species were more abundant in the oral microbiome, which is generally associated with good oral health. However, in PDAC patients, these bacteria were more enriched in the gut. The hypothesis of bacterial translocation where oral bacteria migrate to the gut and contribute to PDAC pathogenesis is not fully supported by these findings. Although shared species were found, the enrichment patterns suggest that these bacteria may have distinct roles in the gut and oral environments of PDAC patients. Further investigation is needed to clarify whether these species actively translocate or if their presence in both microbiomes is due to other underlying factors associated with pancreatic cancer.

When comparing the oral MM panel with the gut MM panel, we found one shared species *Actinomyces SGB17154*. Interestingly, this species was enriched in both the oral and gut microbiomes of male PDAC patients, suggesting its potential involvement in PDAC-associated microbial dysbiosis **(Figure 5C)**. The enrichment of this bacterium in both environments highlights a possible link between the oral and gut microbiomes in PDAC, although further research is needed to fully understand the mechanisms behind these microbiome shifts and their role in pancreatic cancer pathogenesis.

## Discussion

Sex differences in cancer are coming into focus since they may also have relevance for patient outcome ^10^. This study offers significant insights into the sex-specific microbiome dysbiosis in PDAC, highlighting distinct microbial patterns in the oral and gut microbiomes of male and female PDAC patients. The findings support the notion that sex-specific differences in the microbiome may contribute to the heterogeneity in PDAC pathogenesis and response to therapy, and further emphasize the need to incorporate sex as a biological variable in microbiome research.

The study identified notable differences in microbial alpha and beta diversity between PDAC patients and healthy controls, with distinct shifts observed between sexes. Specifically, female PDAC patients showed a significant increase in *Bacteroidetes* and a decrease in *Actinobacteria* in the oral microbiome compared to healthy controls, while male PDAC patients exhibited elevated levels of *Bacteroidetes* and *Proteobacteria*. These phylum-level variations underscore the presence of sex-specific microbial dysbiosis, potentially driven by differences in immune modulation or hormonal influences^11^.

At the genus level, several bacterial genera, including *Prevotella* and *Streptococcus*, displayed differential abundance patterns between male and female PDAC patients, suggesting their possible role in sex-dependent disease progression. *Prevotella*, a genus known for its involvement in inflammatory processes, was significantly enriched in female PDAC patients but not in males, indicating a potential link to sex-specific immune responses in PDAC^12^. Similarly, *Streptococcus*, which has been previously associated with pancreatic cancer, was decreased in both male and female patients, yet the extent of reduction varied between sexes, suggesting different microbial dynamics.

One of the most compelling findings of this study was the identification of shared and sex-specific bacterial species in PDAC patients. Through LEfSe analysis and subsequent Venn diagram comparisons, 25 bacterial species were found to be unique to female patients, while 38 were specific to males, with only 5 species shared between the two groups. This distribution indicates that while some bacterial shifts are common across sexes, the majority of microbiome alterations are sex-specific, likely reflecting underlying differences in host physiology and tumor biology^9^.

Our study also highlighted the potential role of shared bacterial species between the oral and gut microbiomes in PDAC. For female patients, five bacterial species—*Mogibacterium diversum, Streptococcus salivarius, Streptococcus parasanguinis, Rothia mucilaginosa*, and *Streptococcus vestibularis*—were detected in both the oral and gut microbiomes. These species exhibited contrasting enrichment patterns, being more abundant in the oral cavity of healthy individuals and in the gut of PDAC patients, suggesting a possible shift in microbial localization or function in the context of disease. This observation challenges the hypothesis of bacterial translocation as a primary mechanism and instead points towards altered microbial interactions or niche adaptation in PDAC patients^13^. In male PDAC patients, only one species, *Actinomyces* SGB17154, was shared between the oral and gut microbiomes. The consistent enrichment of this bacterium in both environments indicates a stronger microbial link between the two compartments in males, potentially driven by systemic factors such as immune suppression or metabolic changes specific to male PDAC pathogenesis^9^. The limited overlap of shared species between the oral and gut microbiomes further supports the need to investigate sex-specific microbial translocation and its implications for PDAC progression.

Although this study provides valuable insights, there are several limitations. First, the clinical data in the BioProject: PRJNA832909 database lacked detailed information on key factors such as medication, diet, and antibiotic exposure, all of which could influence microbiome composition. Without these variables, it is challenging to draw conclusive associations between microbiome composition and PDAC progression. Future studies should include such factors and employ methods like propensity score matching to control for potential confounders. Second, the relatively small sample size may limit the generalizability of the findings. Future research should involve larger cohorts to increase statistical power, particularly in sex-specific comparisons. Furthermore, the cross-sectional nature of this study makes it difficult to establish a cause-and-effect relationship between microbiome dysbiosis and PDAC. Longitudinal studies will be essential for monitoring microbiome shifts over time and their role in disease development. Third, while the identified bacterial panels for PDAC were sex-specific, further validation in independent cohorts is necessary. Incorporating metabolomic analyses alongside microbiome studies could provide a more comprehensive understanding of how the microbiome contributes to PDAC pathogenesis.

## Conclusion

This study highlights distinct sex-specific microbial patterns in PDAC, demonstrating significant differences in microbial diversity and composition between male and female patients. Using metagenomics, we identified key bacterial species contributing to sex-specific dysbiosis including shared and unique microbial markers in the gut and oral microbiomes. These findings offer opportunities for developing sex-specific diagnostic and therapeutic strategies for PDAC, emphasizing the importance of microbiome analysis in understanding cancer progression.

